# dom2vec: Unsupervised protein domain embeddings capture domains structure and function providing data-driven insights into collocations in domain architectures

**DOI:** 10.1101/2020.03.17.995498

**Authors:** Damianos P. Melidis, Brandon Malone, Wolfgang Nejdl

## Abstract

**Motivation:** Word embedding approaches have revolutionized Natural Language Processing NLP research. These approaches aim to map words to a low-dimensional vector space in which words with similar linguistic features are close in the vector space. These NLP approaches also preserve local linguistic features, such as analogy. Embedding-based approaches have also been developed for proteins. To date, such approaches treat amino acids as words, and proteins are treated as sentences of amino acids. These approaches have been evaluated either qualitatively, via visual inspection of the embedding space, or extrinsically, via performance on a downstream task. However, it is difficult to directly assess the intrinsic quality of the learned embeddings.

**Results:** In this paper, we introduce dom2vec, an approach for learning protein domain embeddings. We also present four *intrinsic* evaluation strategies which directly assess the quality of protein domain embeddings. We leverage the hierarchy relationship of InterPro domains, known secondary structure classes, Enzyme Commission class information, and Gene Ontology annotations in these assessments. These evaluations allow us to assess the quality of learned embeddings independently of a particular downstream task. Importantly, allow us to draw an analog between the local linguistic features in nature languages and the domain structure and function information in domain architectures, thus providing data-driven insights into the context found in the *language* of domain architectures. We also show that dom2vec embeddings outperform, or are comparable with, state-of-the-art approaches on downstream tasks.

**Availability:** The protein domain embeddings vectors and the entire code to reproduce the results are available at https://github.com/damianosmel/dom2vec.

## 1 Introduction

A primary way of how proteins evolve is through rearrangement of their functional/structural units, known as *protein domains* (Moore *et al*., 2008; Forslund and Sonnhammer, 2012). The domains are independent folding and functional modules and so they exhibit conserved sequence segments. Prediction algorithms exploited this information and used as input features the domain composition of a protein for various tasks. For example (Chou and Cai, 2002) classified the cellular location and (Forslund and Sonnhammer, 2008; Doǧan *et al*., 2016) predicted the associated Gene Ontology (GO) terms. There exist two ways to represent domains; either by the linear order in a protein, *domain architecture*, (Scaiewicz and Levitt, 2015), or by a graph where nodes are domains and edges connect domains that co-exist in a protein (Moore *et al*., 2008; Forslund and Sonnhammer, 2012).

Moreover, (Yu *et al*., 2019) investigated if the domains architecture has a grammar as a natural spoken language. They compared the bigram entropy of *domain architecture* for PFAM domains (Sonnhammer *et al*., 1998) to the respective entropy of the English language, showing that although it was lower than the English language, it was significantly different from a language produced after shuffling the domains. Prior to this result, methods had exploited *domain architecture* representation to various applications, such as fast homology search (Terrapon *et al*., 2013) and retrieval of similar proteins (Marchler-Bauer *et al*., 2016).

Word embeddings are unsupervised learning methods which have as input large corpora and they output a dense vector representation of words contained in the sentences of these documents based on the distributional semantic hypothesis, that is the meaning of a word can be understood by its context. Thus a word vector represents local linguistic features, such as lexical or semantical information, of the respective word. Several methods to train word embeddings have been established, for example (Collobert and Weston, 2008; Mikolov *et al*., 2013b; Pennington *et al*., 2014). These representations have been shown to hold several properties such as analogy and grouping of semantically similar words (Mikolov *et al*., 2013a; Drozd et al., 2016). Importantly, these properties are learned without the need of a labeled data set. Word embeddings are currently the mainstream input for neural networks in the Natural Language Processing (NLP) field as first they reduce the feature space (compared to 1-hot representation) and second they provide word features that encapsulate relation between words based on linguistic features. The use of word embeddings improved the performance on most of the tasks (sentiment analysis, Named Entity Recognition (NER), etc).

Various methods to create embeddings for proteins are proposed (Asgari and Mofrad, 2015; Yang *et al*., 2018; Bepler and Berger, 2019; Asgari *et al*., 2019; Heinzinger *et al*., 2019; Alley *et al*., 2019). ProtVec fragmented the protein sequence in 3-mers for all possible starting shifts, then learned embeddings for each 3-mer and represented the respective protein as the average of its constituting 3-mer vectors (Asgari and Mofrad, 2015). SeqVec utilized and extended the Embeddings from Language Models (ELMo) (Peters *et al*., 2018) to learn a dense representation per amino acid residue, resulting in matrix representations of proteins, created by concatenating their learned residue vectors (Heinzinger *et al*., 2019).

Both approaches evaluated the learned representations qualitatively and quantitatively. Qualitatively by having plotted the reduced 2-D embedding space and reported the appearance of cluster of protein with similar properties (biophysical/chemical, structural, enzymatic, taxonomic). For quantitative evaluation, they measured the improvement of performance in down-stream tasks (*protein family classification, cellular location prediction, etc*).

Focusing on their qualitative evaluation, it lacks the ability to assess different learned embeddings in a sophisticated manner. Indeed for word embeddings there is an increase in methods to evaluate words representations intrinsically such as (Schnabel *et al*., 2015). All these techniques are thoroughly summarized by the survey (Lastra-Díaz *et al*., 2019). This is a fruitful direction, because having such evaluation metrics we can validate the knowledge acquired by all trained embedding spaces. And further we can select the best performing space to input to the machine learning algorithm. However creating embeddings for single residues or *k*-mers residues does not enable intrinsic evaluation methods, as biological information for each such fine-grained feature cannot be obtained for all disposed proteins (that embeddings are learned from).

Given that InterPro database (release 70.0) contains domains for 80.9% of the UniProtKnowledgeBase (UniProtKB) sequences (The UniProt Consortium, 2017) and includes associated biological knowledge for the majority of domains (Mitchell and *et al*., 2019), we trained domain embeddings and evaluated them both in intrinsic and extrinsic way.

This work has five major contributions:

- We constructed the *domain architectures* of all UniProtKB sequences with an identified InterPro domain. Then using word2vec method, we learned the vector representation for each InterPro domain based on the constructed *domain architectures*.
- We established intrinsic evaluation methods based on four distinct biological information for a domain. First we evaluated the learned embedding space by domain hierarchy (tree structure capturing family-subfamily relations provided by InterPro). We investigated the performance of a nearest neighbor classifier, 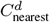, to predict the secondary structure class provided by SCOPe (Fox *et al*., 2013) and the primary EC class in the learned embedding space. We examined the performance of 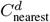 classifier to predict the GO molecular function class for three example model organisms and one human pathogen.
- As a by-product of the intrinsic evaluation, we observed that 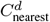 reaches adequate accuracy, compared to 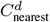 on randomized domains vectors, for secondary structure, enzymatic and GO molecular function. Thus we hypothesized the analog between word embedding clustering by local linguistic features and protein domains clustering by domains structure and function, which in turn, brings data-driven insights into the semantic context of *domain architectures*.
- To evaluate our embeddings extrinsically, we inputted the learned domains embeddings to simple neural networks and compared their performance with state-of-the-art methods in three full-protein tasks; surpassing both state-of-the-art methods for the two tasks, EC primary class and toxin presence, and being comparable in the cellular location prediction task.
- We make available the trained domains embeddings to be used for protein supervised learning tasks by the research community.

## 2 Related work

### 2.1 Protein embedding methods

Works on protein embeddings can be divided in two methods based on their word assignment, the former learn embeddings for constant amino acid subsequences and the latter for variable length subsequences.

#### 2.1.1 Constant length subsequence vectors

Early work in protein embeddings (Asgari and Mofrad, 2015) (ProtVec) belongs to this category. In this work, the authors split the protein sequence in 3-mers with the 3 possible starting shifts. They then used the skip-gram model from (Mikolov *et al*., 2013b) to learn the embeddings for each distinct 3-mer. The embedding for a protein was taken as the mean of all its 3-mers. (Yang *et al*., 2018) adapted ProtVec by using doc2vec (Le and Mikolov, 2014) to find the whole protein embedding. Similarly (Woloszynek *et al*., 2019) computed embeddings for all *k*-mers using a range of k; they also used doc2vec to find the whole protein embedding. (Bepler and Berger, 2019) learned an embedding of each amino acid position incorporating global structural similarity between a pair of proteins and contact map information for each protein. They represented the alignment of two proteins as a soft symmetric alignment of the embeddings of the protein residues. Recently, SeqVec (Heinzinger *et al*., 2019) applied and extended ELMo model (Peters *et al*., 2018) to learn a contextualized embedding per amino acid position. Recently, UniRep was introduced (Alley *et al*., 2019) to learn embeddings for amino acids by a language model that uses RNNs.

#### 2.1.2 Variable-length subsequence vectors

ProtVecX (Asgari *et al*., 2019) embeds proteins by first extracting (variable-length) motifs from proteins using a data compression algorithm, byte-pair encoding (Gage, 1994). ProtVec is then used to learn embeddings for the motifs.

### 2.2 Extrinsic evaluation methods

Extrinsic evaluation methods, like performance on a supervised learning task, are most commonly used to evaluate the quality of embeddings. To date, different papers have evaluated performance on different downstream tasks and using different datasets.

#### Sequence-level prediction

(Asgari and Mofrad, 2015) predicted protein family and disorder, while (Yang *et al*., 2018) evaluated embeddings by predicting the *Channelrhodopsin (ChR)* localization, the thermostability of *Cytochrome P450*, the *Rhodopsin* absorption wavelength and the *Epoxide hydrolase* enantioselectivity. (Bepler and Berger, 2019) predicted transmembrane regions of proteins, and (Heinzinger *et al*., 2019) used subcellular localization and water solubility as validation. (Asgari *et al*., 2019) consider venom toxin, subcellular localization, and enzyme primary class predictions tasks for downstream validation. (Alley *et al*., 2019) predicted stability of proteins and the phenotype of diverse variants of the green fluorescent protein.

#### Per-residue prediction

(Heinzinger *et al*., 2019) predicted secondary structure and intrinsic disorder and (Alley *et al*., 2019) predicted functional effect of single amino acid mutations.

### 2.3 Qualitative evaluation methods

Many of the described approaches use some form of qualitative evaluation. Commonly, the learned embeddings are projected to two dimensions using techniques like tSNE (Maaten and Hinton, 2008) or UMAP (McInnes *et al*., 2018) to visually inspect the embeddings. While such approaches are helpful to gain trust in the embeddings, they do not provide a rigorous approach to compare the quality of multiple embedding spaces, such as those found when using different hyperparameters for the embedding models.

(Asgari and Mofrad, 2015) used a qualitative intrinsic evaluation approach for ProtVec. They visualized the 2-dimensional reduced protein space along with their biophysical and biochemical properties to report the appearance of clusters with similar biochemical properties.

For intrinsic evaluation they (Heinzinger *et al*., 2019) visualized the 2-dimensional reduced protein space along with secondary structure class, the EC and the taxonomic kingdoms per protein.

## 3 Materials and methods

Our approach is summarized in Figure 1, in the following we explain the methodology for each part of the approach.

**Fig. 1.**
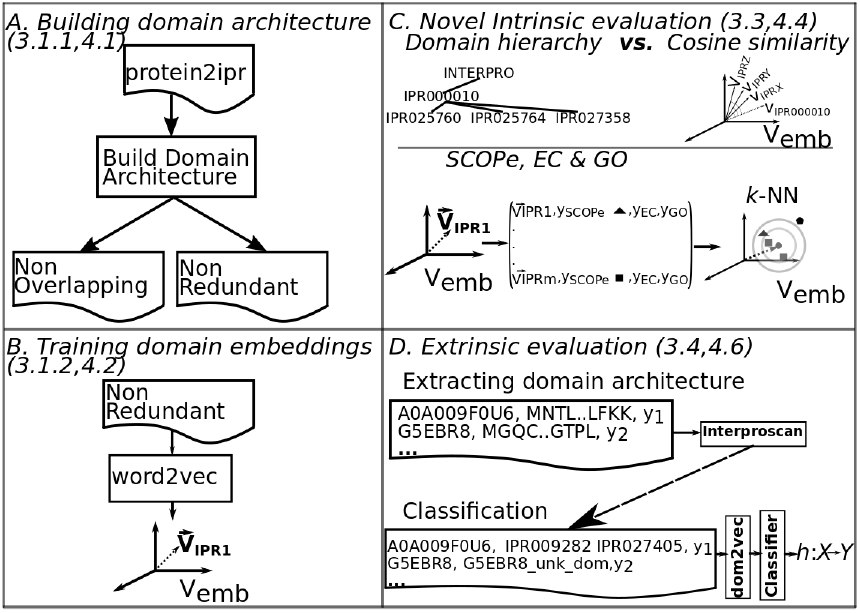
Summary of our approach divided in four parts, building two forms of domains architecture, training domain embeddings, performing intrinsic and extrinsic evaluation of embeddings. Numbers in parentheses indicate the sections discussing the respective part.

### 3.1 dom2vec

We now describe dom2vec, our novel approach for learning protein domain embeddings based on domain architectures.

#### 3.1.1 Building domain architecture

Hereafter we will refer to the sequence of domains in a protein as its *domain architecture*. We consider two distinct strategies to represent a protein based on its *domain architecture: non-overlapping* and *non-redundant*. In both cases, the sequences are based on protein domain entries from Interpro. For efficiency, an interval tree is built for each protein to detect overlapping domains, and each protein is split into regions with overlapping domains.

##### Non-overlapping sequences

For each region with overlapping domains, all domains except the longest are removed.

##### Non-redundant sequences

For each region with overlapping domains, all domains with the same Interpro identifier, except the longest domain with each identifier, are removed.

In both cases, the domains in each protein are sorted by start position to construct its *domain architecture*. Following the approach of (Doǧan *et al*., 2016) we also added the “GAP” domain to annotate more than 30 amino acid subsequence that does not match any InterPro domain entry.

#### 3.1.2 Training domain embeddings

Next, we learned task-independent embeddings for each domain using two variants of word2vec (Mikolov *et al*., 2013b): continuous bag of words CBOW and SKIP. See (Mikolov *et al*., 2013b) for technical details on the difference between these approaches. In our context, each domain is taken as a word, and each protein sequence is considered as a sentence. Thus, after learning, each domain is associated with a task-independent embedding.

In Section 3.3, we consider several *intrinsic* evaluation strategies to determine the quality of these embeddings for various hyperparamers of word2vec.

### 3.2 Qualitative evaluation

As a preliminary evaluation strategy, we used qualitative evaluation approaches adopted in existing work. To follow the qualitative approach of ProtVec and SeqVec we also visualized the embedding space for selected domain superfamilies, to answer the following research question,

*RQ:* *Did vectors of each domain superfamily form a cluster in the V_emb_?*

#### Evaluation

To find out, we added the vector of each domain in a randomly chosen domain superfamily to an empty space. Then we performed principle component analysis (PCA) (Pearson, 1901) to reduce the space in two dimensions and observed the formed clusters.

### 3.3 Novel intrinsic evaluation methods

Previous work has evaluated the quality of embeddings only indirectly by measuring performance on downstream, supervised tasks, as described in Section 3.4. However, in the natural language processing field, the quality of a learned word embedding space is often evaluated *intrinsically* by considering relationships among words, such as analogies. Such an evaluation is important because it ensures the learned embeddings are meaningful without choosing a specific downstream task.

Such intrinsic evaluations cannot be performed for the ProtVec and SeqVec protein embeddings, because in contrast with these approaches dom2vec directly learns embeddings for protein domains. That is, protein domains are typically represented using structures like profile hidden Markov models (HMM) (Eddy, 1998) rather than unique amino acid sequences. Thus it is not straightforward how sequence feature embeddings can be combined for a given profile HMM in order to evaluate the learned sequence embeddings in the way that we will describe next.

Thus, prior approaches, such as ProtVec, SeqVec, *cannot* learn unique embeddings per domain not allowing us to perform the same evaluation tasks for the existing protein embeddings.

We propose four intrinsic evaluation approaches for domain embeddings: domain hierarchy based on the family/subfamily relation, secondary structure class, Enzymatic Commission (EC) primary class, and Gene Ontology (GO) molecular function annotation.

We refer to the embedding space learned by word2vec for a particular set of hyperparameters as *V_emb_*. We refer to the *k* nearest neighbors of a domain *d* as 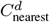 found using the Euclidean distance. To inspect the relative performance of *V_emb_* on each of the following evaluations, we *randomized* all domain vectors and run each evaluation task. That is, irrespective of the embedding method parameters and only for each different space dimensionality we assigned to each domain vector a newly created random vector and save this random space to be evaluated.

#### 3.3.1 Domain hierarchy

InterPro defines a strict family-subfamily relationship among domains. This relationship is based on sequence similarity of the domain signatures. We refer to the children of domain *p* as *C^p^*. We use these relationships to evaluate an embedding space, posing the following research question,

*RQ:* *Did vectors of hierarchically closely domains form clusters in the V_emb_?*

##### Evaluation

We evaluate *V_emb_* by retrieving 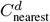 of each domain. For all learned embedding spaces, we measured their recall performance, *Recall_hier_* defined as follows:

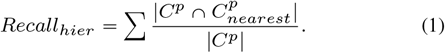

#### 3.3.2 Secondary structure class

We extracted the secondary structure of Interpro domains from the SCOPe database and form the following research question,

*RQ:* *Did vectors of domains, with same secondary structure class, form clusters in the V_emb_?*

##### Evaluation

We evaluated *V_emb_* by retrieving 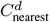 of each domain. Then we applied stratified cross-fold validation and measured the performance of a *k*-nearest neighbor classifier to predict the structure class of each domain. The intrinsic evaluation performance metric is the average accuracy across all folds, *Accuracy_SCOPe_*.

#### 3.3.3 Enzyme Commission (EC) primary class

The enzymatic activity of each domain is given by its primary EC class (Fleischmann *et al*., 2004) and pose the following research question,

*RQ:* *Did vectors of domains, with same secondary structure class, form clusters in the V_emb_?*

##### Evaluation

We again evaluate *V_emb_* using *k* nearest neighbors in a stratified cross-validation setting. Average accuracy across all folds is again used to quantify the intrinsic quality of the embedding space.

#### 3.3.4 GO molecular function

For our last intrinsic evaluation, we aimed to assess *V_emb_* using the molecular function GO annotation. We extracted all molecular function GO annotations associated with each domain.^1^ Since model organisms have the most annotated proteins we created GO molecular function data sets for one example prokaryote (*Escherichia coli* denoted *E.coli*), one example simple eukaryote (*Saccharomyces cerevisiae* denoted *S.cerevisiae*) and one complex eukaryote (*Homo sapiens* denoted *Human*). To assess our embeddings also for not highly annotated organisms, we included a molecular function data set for an example human pathogen (*Plasmodium falciparum*, denoted as *Malaria*). Finally, we pose the following research question,

*RQ:* *Did vectors of domains, with same GO molecular function, form clusters in the V_emb_?*

##### Evaluation

We again evaluate an embedding space using *k* nearest neighbors in a stratified cross-validation setting. Average accuracy across all folds is again used to quantify performance.

### 3.4 Extrinsic evaluation

In addition to assess the learned *V_emb_* we also examine the performance change in downstream tasks. That is for three data sets-supervised tasks, we feeded the domain representations in simple neural networks and compare the performance of our model with state-of-the-art protein embeddings.

#### 3.4.1 Simple neural models for downstream tasks

We consider a set of simple, well-established neural models to combine the domain embeddings for each protein to perform downstream tasks, that is, for *extrinsic* evaluation tasks. In particular, we use FastText (Joulin *et al*., 2017), convolutional neural networks (CNNs) (LeCun *et al*., 1998), and recurrent neural networks (RNNs) with long-short term memory (LSTM) cells (Hochreiter and Schmidhuber, 1997) and bi-directional LSTMs. We leave evaluation with more sophisticated models, such as transformers like BERT (Devlin *et al*., 2018), for future work.

The location data set is a multi-class task, thus we used as performance metric a multi-class generalization of area under the receiver operating characteristic curve (mc-AuROC). For the membrane data set which is a binary task, the performance metric was the area under the receiver operating characteristic curve (AuROC).

#### 3.4.2 TargetP

We downloaded the TargetP data set provided by (Emanuelsson *et al*., 2000). To compare with ProtVecX we also used the non-plant data set. This data set consists of 2,738 proteins accompanying with their uniprot id, sequence and the cellular location label which can be nuclear, cytosol, pathway or signal and mitochondrial. Finally, we removed all instances with duplicate set of domains, resulting in total of 2,418. This is a multiclass task and its class distribution is summarized in supplementary section E.

##### Evaluation

For the TargetP we used the mc-AuROC performace metric.

#### 3.4.3 Toxin

(Gacesa *et al*., 2016) introduced a data set associating protein sequence to toxic or other physiological content. To compare with ProtVecX we used the hard setting which provides uniprot id, sequence and the label toxin content or non-toxin content, for 15,496 proteins. Finally, we kept only the proteins with unique domain composition, resulting to 2,270 protein instances in total. This is a binary task and the class distribution is shown in supplementary section E.

##### Evaluation

As Toxin data set is binary task, we used AuROC as performance metric.

#### 3.4.4 NEW

We downloaded the NEW data set from (Li *et al*., 2017). For each of the 22,618 proteins the data set provides sequence and the EC number (class). The primary enzyme class, first digit of EC number, is our label on this prediction task, resulting in a multi-class task. Finally, we removed all instances with duplicate domain composition, resulting in a total of 14,434 protein instances. The possible classes are 6 and the class distribution is shown in supplementary section E.

##### Evaluation

NEW data set is a multi-class task, thus we used mc-AuROC as performance metric.

#### 3.4.5 Data partitioning

We divided each data set into 70/30% train and test splits. To perform model selection, we created inner three-fold cross validation sets on each train split.

Then we observed that the performance of classifier depending on protein domains is highly dependent on the out-of-vocabulary (OOV) domains, as first discussed in (Luong *et al*., 2015). OOV domains are all the domains contained in the test set, *but* not in the train. For TargetP, Toxin and NEW we observed that approximately 60%, 20%, 20% of test proteins contain *at least one* OOV domain. Thus, we run experiments based on this observation. For TargetP, containing the highest OOV, we split the test set into shorter sets by an increasing degree of OOV, namely 0%, 10%, 30%, 50%, 70%, 100%. Then we learned models for the whole train set and benchmarked the performance on each of these test subsets.

For Toxin and NEW data set, experiencing low OOV, we seek to investigate the generalization of the produced classifier, by increasing the number of training examples that the model learn from and benchmark always in the entire test set. Thus, we created training splits of size, 10%, 20%, 50% of the whole train set. To perform significance testing we created 10 random subsamples for each training split percentage.

## 4 Results and discussion

### 4.1 Building domain architecture

We downloaded the domain hits, protein2ipr.dat, for UniProt proteins from the InterPro version 75. This file contained 128 660 257 proteins with InterPro signature, making up the 80.9% of the total UniProtKB proteome (version 2019_06). For all these proteins we extracted the non-overlapping and non-redundant sequences, which we processed in the next section. The number of unique non-overlapping sequence was (35 183 + 1) plus the “GAP” domain and non-redundant was (36 872 + 1) plus the “GAP”. Comparing to total number of domains in InterPro version 75, which was 36 872, we observed that non-overlapping sequences captured 95.42% and the non-redundant captured 100% of the InterPro domains. Figure 2 demonstrates an example of constructed domain architectures by showing the domain architectures for the subdomains for the Tetrapyrrole methylase domain of the *Diphthine synthase* protein (see (Mitchell and *et al*., 2019) for more details).

**Fig. 2.**
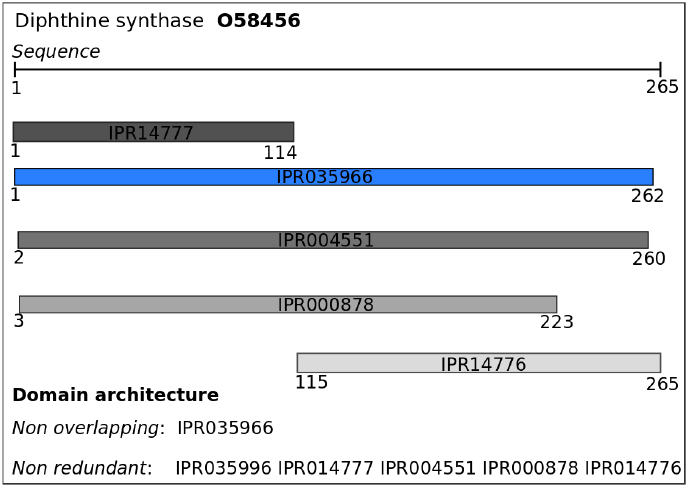
Non overlapping and non-redundant domain architecture of Diphthine synthase protein. Because all domains are overlapping with the largest one, colored in blue, the nonoverlapping sequence is just the single longest domain (IPR035966). All other domains have a unique InterPro id, so the set of non-redundant sequences includes all presented domains sorted by starting position; we colored all these other domains in fading hues of gray based on their starting position.

### 4.2 Training domain embeddings

#### 4.2.1 Domain architecture

Before we applied the word2vec package we examined the histograms of number of domains per protein for non-overlapping and non-redundant sequences. We show the histogram of the number of non-overlapping sequences per protein in Figure 3. We observed that these distributions are skewed and long-tailed. Then we used both CBOW and SKIP algorithms to learn domain embeddings. We used the following parameters sets. Based on the histograms we picked the window parameter for word to be 2 or 5, *w* = {2, 5}. For the number of dimensions, we used common values from the NLP literature, *dim* = {50,100, 200}. We trained the embeddings from 5 to 50 epochs with step size 5 epochs *ep* = {5,10,15,…, 50}. Finally, all other parameters were set to their default values. For example, the negative sampling parameter was left to default, *ng*=5.

**Fig. 3.**
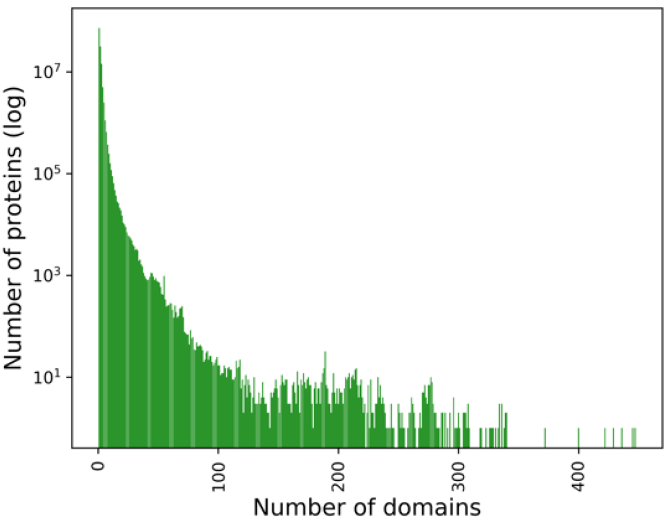
Histogram of non-overlapping domains per protein, where the number of proteins is in log_10_ scale.

### 4.3 Qualitative evaluation

We randomly selected five InterPro domain superfamilies to perform the experiment. The selected domain superfamilies were *PMP-22/EMP/MP20/Claudin superfamily* with parent InterPro id IPR004031, *small GTPase superfamily* with parent InterPro id IPR006689, *Kinase-pyrophosphorylase* with parent InterPro id IPR005177, *Exonuclease, RNase T/DNA polymerase III* with parent InterPro id IPR013520 and *SH2 domain* with parent InterPro id IPR000980. Similarly to domain hierarchy, we loaded the parent-child tree *T_hier_*, provided by InterPro, and for each domain superfamily starting from the parent domain we included recursively all domains that have subfamily relation with this parent domain. For example the *Kinase-pyrophosphorylase* domain superfamily has domain parent IPR005177, which in turn has two immediate domain subfamilies IPR026530, IPR026565 and IPR026565 has domain subfamily IPR017409, consequently the set of domains for *Kinase-pyrophosphorylase* domain superfamily is {IPR005177,IPR026530,IPR026565,IPR017409}.

We retrieved the vectors for each domain in a domain superfamily using the 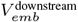, the best performing dom2vec space selected in Section 4.5. Finally, we visualized the two-dimensional PCA-reduced space at Figure 4. From this Figure 4, we observed that domains embeddings of each superfamily form well-separated clusters; the cluster of *Exonuclease, RNase T/DNA polymerase III* superfamily has the highest dispersion of all presented superfamilies.

**Fig. 4.**
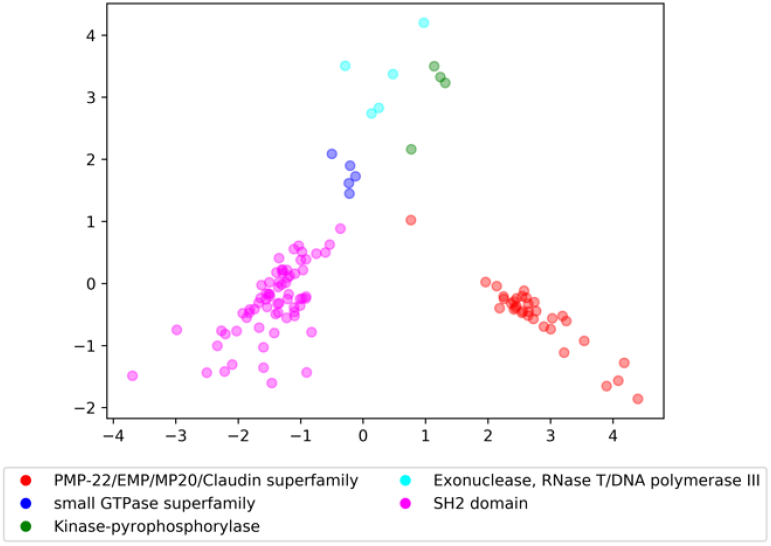
Visualization of domain vectors for five domain superfamilies in best performing dom2vec space 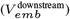.

### 4.4 Novel intrinsic evaluation

In the following, we evaluated each instance of learned embedding space *V_emb_* for both non-overlapping and non-redundant representations of domain architectures. An instance of *V_emb_* space is the embeddings space learned for each combination of the product *non_overlap* × *w* × *dim* × *ep*. Thus, the total number of embeddings spacesis |*non_overlap*| × |*w*| × |*dim*| × |*ep*| = 2×2×3×10 = 120. Let 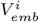 denote such embedding space instance. In the following subsection we evaluated each 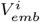 instance for domain hierarchy, secondary structure, enzymatic process and GO molecular function.

#### 4.4.1 Domain hierarchy

We loaded the parent-child tree *T_hier_*, provided by InterPro, consisting of 2 430 parent domains. Then for each 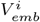 we compared the actual and predicted children of each parent, and we averaged out the recall for all parents. For ease of presentation, we show only the results for non-redundant sequences at Table 1 and we provide the complete results in the Supplementary A. However, in the following we discuss the results.

**Table 1.** Average *Recall_hier_* for non-redundant sequences, best shown in bold case

| Emb. Meth, $w$ | $dim=50$ | $dim=100$ | $dim=200$ |
| --- | --- | --- | --- |
| CBOW, $w=2$ | 0.406 ($ep=10$) | 0.412 ($ep=10$) | 0.414 ($ep=5$) |
| CBOW, $w=5$ | 0.405 ($ep=30$) | 0.402 ($ep=35$) | 0.382 ($ep=10$) |
| SKIP, $w=2$ | 0.512 ($ep=5$) | 0.53 ($ep=5$) | **0.538** ($ep=5$) |
| SKIP, $w=5$ | 0.507 ($ep=5$) | 0.525 ($ep=5$) | 0.524 ($ep=5$) |
| random | 0 | 0 | 0 |

From Tables S1 and 1 (appendix A), we observed that SKIP performed better overall, and the embeddings learned from non-redundant annotations always have better average recall values compared to nonoverlapping. The best performing 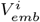 achieved average *Recall_hier_* of 0.538; that embedding space was trained using non-redundant annotations with SKIP, *w* = 2 and *dim* = 200. We plotted the histogram of recall values for this best performing space, Figure S1 (Suppl. A), and we observed that the embeddings space brought close domains with *not known* family-subfamily relation for almost the one third of the parent domains (827 out of 2 430). To diagnose the reason for this moderate performance, we plotted the histogram of the number of children for each parent having recall 0, Figure S2 (Suppl. A). We observed that most of these parents have only one child. Consequently, the embedding space should have been very precise for each of these parent child relation in order to acquire better recall than 0. We compared this moderate performance of *V_emb_* with the performance of the randomized spaces, which was equal to 0. Thus, we concluded that our embedding spaces greatly outperformed each randomized space for domain hierarchy relation. Consequently, we can answer our initial question: the majority of domains of the same hierarchy are placed in close proximity in the embedding space.

#### 4.4.2 Secondary structure class

We extracted the SCOPe class for each InterPro domain from the interpro.xml file. This resulted to 25 196 domains with unknown secondary structure class, 9 411 with single secondary structure class and 2 265 domains with more than one assigned classes (multi-label). For clarity, we removed all multi-label and unknown instances leaving with 9 411 single-labeled instances; the class distribution of the resulting data set is shown in Table S2 (Supplementary B).

To answer the research question, we measured the performance of 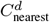 classifier in each 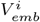 to examine the homogeneity of the space with respect to the SCOPe class. We split the 9 411 domains in 5-fold stratified cross validation sets. To test the change in prediction accuracy for an increasing number of neighbors, we used different sets of neighbors, namely, *k* = {2, 5, 20, 40}. We summarized the results for the best performing 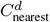 which was for *k* = 2 for non-redundant sequences in 2. We show the respective table for non-overlapping sequences in the Suppl. B. We compared these accuracy measurements to the respective ones of the random spaces, and we found that the lowest accuracy values, achieved for CBOW *w*=5 using non-overlapping domains, are twice as high as the accuracy values of the random spaces for all possible dimensions. We observed again that the non-redundant domains resulted in higher accuracy compared to non-overlapping domains. The best performing 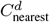 was for 2-NN for (non-redundant, SKIP, *w*=5, *dim*=50, *ep*=25) with average accuracy over the folds equal to 84.56. Consequently, we can answer our second research question: domain embeddings of the same secondary structure class form distinct clusters in a learned embedding space.

#### 4.4.3 Enzyme Commission (EC) primary class

To this end we extracted the EC primary class from the interpro.xml file. This resulted in 29 354 domains with unknown EC, 7 248 domains with only one EC and 721 with more than one EC. As before, we removed all multi-label and unknown instances leaving 7 428 domains with known EC. For each 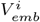 we augmented a domain instance with its vector representation, and then we used 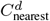 to predict the EC. Please refer to the Supplementary C for the class distribution of EC task. We again omit the entire result tables and we discuss the main results.

We present the average *Accuracy_EC_* obtained on embedding space learned using non-redundant sequences in Table 3. We show the respective table for non-overlapping in the Suppl. C. We compared these accuracy measurements to the respective ones of the random spaces. We found that the minimum average *Accuracy_EC_* value was equal to 60.51 and was achieved using CBOW *w*=5 for non-overlapping sequences. That value is approximately twice as large than the accuracy values of the random spaces for all possible dimensions (maximum average *Accuracy_EC_* for random space with *dim*=100 was 32.64). The best results were always produced using 2-NN, and we saw once more that SKIP trained on non-redundant sequences performed the best with maximum average accuracy of 90.85 (SKIP, *w*=5, *dim*=50, *ep*=30). Consequently, we can answer our third research question: domain embeddings of the same EC primary class form distinct clusters in a learned embedding space.

**Table 3.**
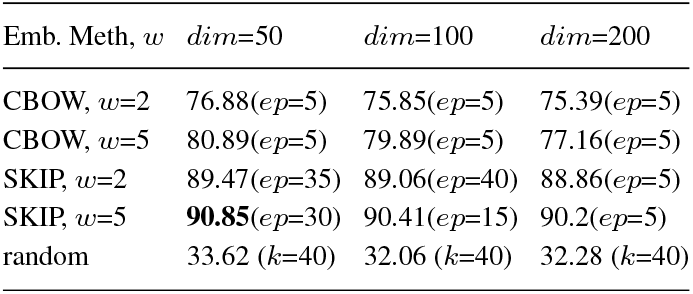
Average 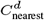 *Accuracy_EC_* non-redundant sequences (*k*=2), best shown in bold case

| Emb. Meth, $w$ | $dim=50$ | $dim=100$ | $dim=200$ |
| --- | --- | --- | --- |
| CBOW, $w=2$ | 76.88($ep=5$) | 75.85($ep=5$) | 75.39($ep=5$) |
| CBOW, $w=5$ | 80.89($ep=5$) | 79.89($ep=5$) | 77.16($ep=5$) |
| SKIP, $w=2$ | 89.47($ep=35$) | 89.06($ep=40$) | 88.86($ep=5$) |
| SKIP, $w=5$ | **90.85**($ep=30$) | 90.41($ep=15$) | 90.2($ep=5$) |
| random | 33.62 ($k=40$) | 32.06 ($k=40$) | 32.28 ($k=40$) |

#### 4.4.4 GO molecular function

We parsed the GO annotation file of InterPro to extract first-level GO molecular function for domains for the four organisms. We followed the same methodology to examine the homogeneity of a *V_emb_* with respect to GO molecular function annotations. Thus, for each 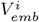, we augmented each domain by its vector and its GO label, and we classified each domain using 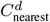. As before, we used 5-fold stratified cross-validation for evaluation. In our experiments, we varied the number of neighbors *k* = {2, 5, 20, 40} to test its influence to the change of performance. For space limitations, we summarized the performances showing only the best average accuracy over the number of neighbors. For the ease of presentation we omit the resulted tables for three first organisms and show only for *Human*; however, we discuss the results for all organisms. Please see supplementary D for full results.

**Table 2.**
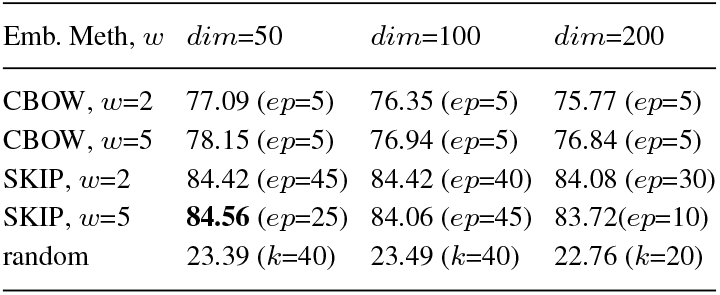
Average 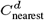 *Accuracy_SCOPe_* for non-redundant sequences (*k*=2), best shown in bold case

For *Malaria*, the best average accuracy was 76.86, scored by the 2-NN for the 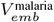 corresponding to (non-redundant, SKIP, *w*=5, *dim*=100, *ep*=40). The minimum accuracy value was 56.94 and was achieved using CBOW with *w*=5 using non-overlapping sequences. We compared this minimum accuracy to the maximum accuracy obtained in a random space, which was 47.57. That is, even the worst-performing dom2vec approach outperformed the random baseline by ten percent.

For *E.coli*, the best average accuracy was 81.72, scored using 2-NN for the 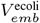 corresponding to (non-redundant, SKIP, *w*=5, *dim*=50, *ep*=5). The minimum accuracy value was 67.34, achieved using CBOW with *w*=5 using non-overlapping sequences. For *E.coli*, the random baseline was still worse, with an accuracy of 64.46.

For *Yeast*, the best average accuracy was 75.10, scored using 5-NN for the 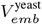 corresponding to (non-redundant, SKIP, *w*=5, *dim*=50, *ep*=50). The minimum accuracy value was 59.82, achieved using CBOW with *w*=5 using non-overlapping sequences. We compared this miminum accuracy to the maximum accuracy obtained in a random space, which was 53.73.

For *Human*, the best average performance for non-redundant sequences are shown in Table 4. The best average accuracy was 75.96, scored by 2-NN for the 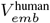(non-redundant, SKIP, *w*=5, *dim*=50, *ep*=40). We compared the minimum accuracy values, obtained by CBOW with *w*=5, to accuracy values of the random spaces, and we found that the worst-performing dom2vec was 20 percentage values higher than the random baseline.

**Table 4.** Average 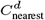 *Accuracy_GO_* for non-redundant sequences, when *k* is not shown *k*=2, best shown in bold case

| Emb. Meth, $w$ | $dim=50$ | $dim=100$ | $dim=200$ |
| --- | --- | --- | --- |
| CBOW, $w=2$ | 66.94($ep=5$) | 66.32($ep=5$) | 66.32($ep=5$) |
| CBOW, $w=5$ | 67.77($ep=5$) | 65.87($ep=5$) | 65.77($ep=5$) |
| SKIP, $w=2$ | 74.77($ep=40$) | 74.18($ep=5$) | 73.14($ep=5$) |
| SKIP, $w=5$ | **75.96**($ep=40$) | 75.53($ep=10$) | 74.98($ep=5$) |
| random | 37.05 ($k=40$) | 37.03 ($k=20$) | 37.05 ($k=40$) |

For all four example organisms, we observed that the SKIP on non-redundant sequences produced *V_emb_* where 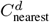 achieved the best average accuracy in. For the three out of the four organisms, the best performances were achieved for the lowest number of dimensions (*dim*=50). In all cases, we found that the worst-performing dom2vec embeddings outperformed the random baselines.

### 4.5 Selecting trained embeddings

Based on the previous four experiments, we aimed to evaluate the learned *V_emb_* spaces and select the best domain embedding space for downstream tasks. In all experiments, the SKIP algorithm for non-redundant sequences created best performing embedding spaces. From the individual results, we saw that the configuration of parameters (non-redundant, SKIP, *w*=5, *dim*=50) brought best results in 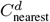 performance for SCOPe, EC and GO for *E.coli, Yeast, Human*, second best for *Malaria* and the sixth best average recall (0.507) for the domain hierarchy relation. Therefore, for the next of the work 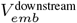 denotes the space produced by (non-redundant, SKIP, *w*=5, *dim*=50).

### 4.6 Extrinsic evaluation

#### 4.6.1 Extracting domain architecture

For each data set that contained the UniProt identifier for the protein instance, we extracted the domain architecture for non-redundant sequences, already created in Section 4.1. For all proteins whose UniProt identifier could not be matched, or for data sets not providing the protein identifier, we use InterProScan (Jones and *et al*., 2014) to find the domain hits per protein. For proteins without a domain hit after InterProScan, we created a protein-specific, artificial protein-long domain; for example, for the protein G5EBR8, we assigned a protein-long domain named “G5EBR8_unk_dom”.

#### 4.6.2 Model selection

To select which simple neural model we should compare to the baselines, we perform hyperparameter selection using an inner, three-fold cross validation on the training set; the test set is not used to select hyperparameters. We used common parameters, dropout of 0.5, batch size of 64, the Adam optimizer (Kingma and Ba, 2015) with learning rate of 0.0003, weight decay of 0 and number of epochs equal to 300. As a final hyperparameter, we allowed updates to the learned domain embeddings, initialized by 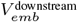. The results are shown in Supplementary E.

### 4.7 Running baselines

Then, we used the same network as the one in right side of Figure 5 of (Heinzinger *et al*., 2019); we refer to this network as SeqVecNet. Namely, the network first averages the 100 (ProtVec) or 1 024 (SeqVec) dimensional embedding vector for a protein; it then applies a fully connected layer to compress a batch of such vectors into 32 dimensions. Next, a ReLU activation function (with 0.25 dropout) was applied to that vector, followed by batch normalization. Finally, another fully connected layer was followed by the prediction layer. As third baseline, we added the *1-hot* of domains in order to investigate the performance change compared to dom2vec learned embeddings.

**Fig. 5.**
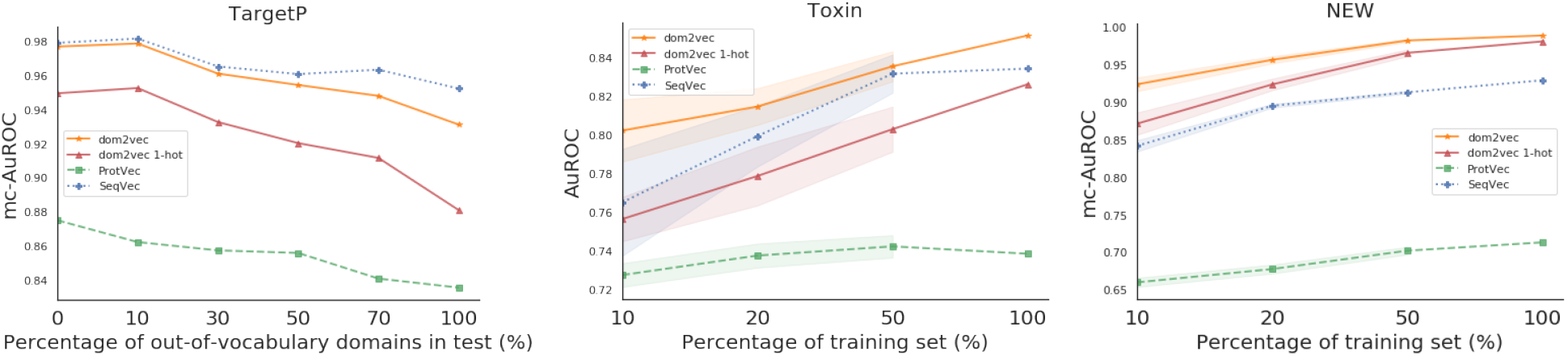
TargetP, OOV experiment: learning in whole train and benchmark in test splits of increasing out-of-vocabulary degree. Toxin and NEW, generalization experiment: learning in increasing train splits, 10 replicates each, and benchmark in whole test sets. The marked points represent the mean performance on the test set and the shaded regions show one standard deviation above and below the mean.

#### 4.7.1 Evaluation

As discussed in 3.4.5, we performed two kinds of experiments. For TargetP, we sought to investigate the effect of OOV on the produced classifier compared to sequence-based embeddings classifiers, which do not experience OOV as their used sequence features are highly common in both train and test set. For Toxin and NEW datasets, we benchmarked the generalization of the produced classifier compared to the sequencebased embeddings classifiers. Finally, for both kinds of experiments, we used the trained models on each test set. Thus, this evaluation shows how differences in the training set affect performance on the test set. The produced performances are shown in Figure 5.

#### 4.7.2 Statistical significance

For both Toxin and NEW, dom2vec significantly outperformed SeqVec, ProtVec and domains *1-hot* vectors. In Toxin data set, we observed that ProtVec learned the less variant model, but with the trade-off obtaining the lowest performance (mc-AuROC). For NEW data set, the *1-hot* representation was the second representation outperforming SeqVec and ProtVec allowing us to validate the finding that domain composition is the most important feature for enzymatic function prediction (Li *et al*., 2017).

For TargetP we validated our assumption that OOV will affect the performance of domains dependent classifiers. That is, for OOV in the range of 0 – 30% the dom2vec classifier was comparable to the best performing model, SeqVec. However, when OOV increased more, then the performance of our model dropped, but still being competitive with the SeqVec. OOV is an known problem in NLP and as future work we would like to investigate ways to resolve this issue. Finally, dom2vec greatly outperformed the 1 – *hot* representation, validating the NLP assumption that unsupervised embeddings improve classification on unseen words, in this context protein domains, compared to 1 – *hot* word (domain) vectors.

## 5 Conclusions

We presented dom2vec, an approach for learning protein domain embeddings using the *domain architectures* deposed in InterPro.

We introduced a novel intrinsic evaluation based on the four sources of biological knowledge of protein domains. We found that the dom2vec vectors cluster by secondary structure, enzymatic function and GO molecular function however such representation could not capture domain hierarchy mostly for domain families of low cardinality. This result comes along with similar results in word embeddings, where it has been shown that, word vectors cluster by semantic and lexical similarity. This byproduct, allows us to draw an analog between words in natural languages and protein domains. Further this finding supports, in a data-driven way, the accepted modular evolution of proteins (Moore *et al*., 2008) and enhance the grammar of protein domain architectures (Yu *et al*., 2019) by giving insights on “forms” of collocations that can be found in such grammar.

In downstream task evaluation, dom2vec outperformed significantly domain 1 – *hotvectors* and state-of-the-art sequence-based embeddings for Toxin and NEW data sets. For the TargetP dom2vec was comparable to the best performing sequence-based embedding, Seqvec, for OOV up to 30%. Thus we recommend to use dom2vec for prediction experiments when the OOV is limited, otherwise sequence-based embeddings can be used.

Finally, we hope that these embeddings will be used for prediction tasks, as well as, for creating data-driven hypothesis to augment our understanding of protein domain architectures.

## Supplementary Information

### Supplementary A: Domain hierarchy

Average recall for all InterPro parents in *T_hier_*, see main paper, for no overlapping sequences are shown in Tables S1. The histogram of average recall for best performing embedding space is shown at Figure S1. Finally, as a first diagnostic, the histogram of the number of children for each parent having *Recall_hier_*=0, is shown at Figure S2.

**Fig. S1.**
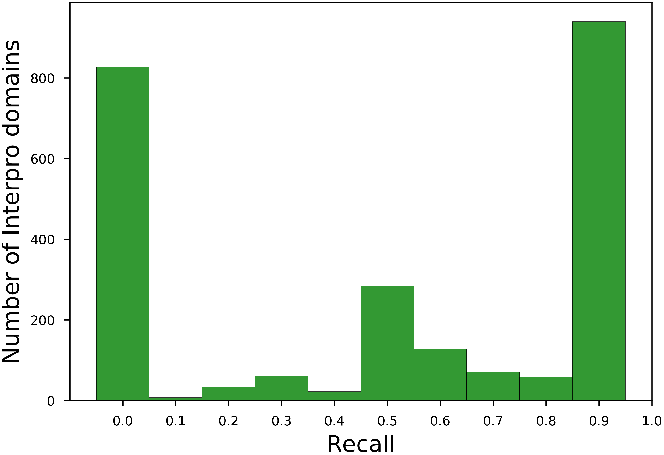
*Recall_hier_* histogram for SKIP, *w*=2, *dim*=200, *ep*=5 for non-redundant annotations.

**Fig. S2.**
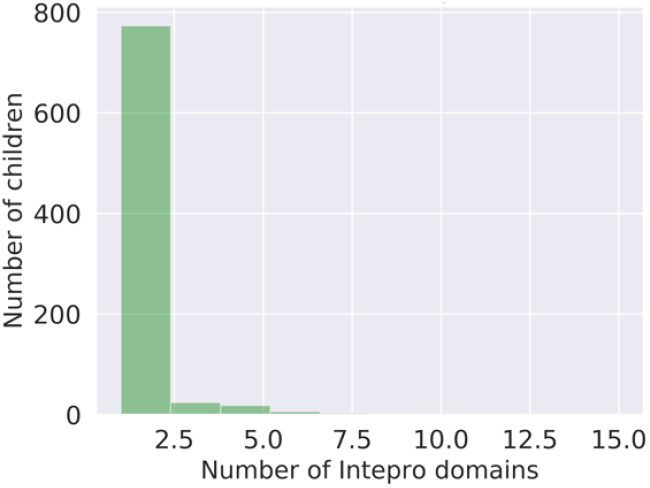
Histogram of number of children for parents with *Recall_hier_*=0.

**Table S1.** Average *Recall_hier_* for non-overlapping sequences, best shown in bold case

| Emb. Meth, $w$ | $dim=50$ | $dim=100$ | $dim=200$ |
| --- | --- | --- | --- |
| CBOW, $w=2$ | 0.242 ($ep=15$) | 0.259 ($ep=20$) | 0.263 ($ep=15$) |
| CBOW, $w=5$ | 0.242 ($ep=45$) | 0.252 ($ep=30$) | 0.25 ($ep=15$) |
| SKIP, $w=2$ | 0.287 ($ep=20$) | **0.316** ($ep=30$) | 0.32 ($ep=20$) |
| SKIP, $w=5$ | 0.284 ($ep=20$) | 0.302 ($ep=30$) | 0.311 ($ep=30$) |
| random | 0 | 0 | 0 |

### Supplementary B: Secondary structure class

Classes distribution of secondary structure class is shown at Table S2. Average 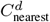 *Accuracy_SCOPe_* for non-overlapping sequences shown in Tables S3.

**Table S2.** SCOPe class summary

| SCOPE class | No. of domains |
| --- | --- |
| a | 1,868 |
| b | 1,806 |
| alb | 2,303 |
| a+b | 2,320 |
| multi-domain | 304 |
| membrane/cell | 309 |
| small | 501 |

**Table S3.** Average 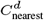 *Accuracy_SCOPe_* for nonoverlapping sequences (*k*=2), best shown in bold case

| Emb. Meth, $w$ | $dim=50$ | $dim=100$ | $dim=200$ |
| --- | --- | --- | --- |
| CBOW, $w=2$ | 50.01($ep=30$) | 50.69($ep=25$) | 50.45($ep=20$) |
| CBOW, $w=5$ | 49.59($ep=25$) | 50.03($ep=25$) | 48.82($ep=15$) |
| SKIP, $w=2$ | **51.83**($ep=30$) | 51.79($ep=20$) | 51.78($ep=15$) |
| SKIP, $w=5$ | 51.54($ep=35$) | 51.65($ep=15$) | 51.34($ep=15$) |
| random | 22.75 ($k=40$) | 24.18 ($k=40$) | 23.39 ($k=40$) |

### Supplementary C: Enzyme Commission (EC) primary class

Classes distribution of EC primary class is shown at Table S4. Average 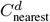 *Accuracy_EC_* for non-overlapping and non-redundant sequences shown in Tables S5 and 3 respectively.

**Table S4.**
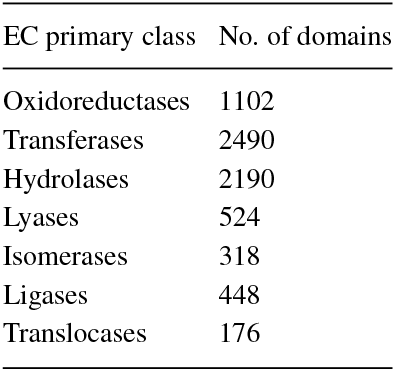
EC class summary

| EC primary class | No. of domains |
| --- | --- |
| Oxidoreductases | 1102 |
| Transferases | 2490 |
| Hydrolases | 2190 |
| Lyases | 524 |
| Isomerases | 318 |
| Ligases | 448 |
| Translocases | 176 |

**Table S5.** Average 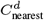 *Accuracy_EC_* non-overlapping sequences (*k*=2), best shown in bold case

| Emb. Meth, $w$ | $dim=50$ | $dim=100$ | $dim=200$ |
| --- | --- | --- | --- |
| CBOW, $w=2$ | 61.23($ep=10$) | 61.33($ep=10$) | 60.66($ep=15$) |
| CBOW, $w=5$ | 61.22($ep=20$) | 60.51($ep=10$) | 60.61($ep=15$) |
| SKIP, $w=2$ | 63.56($ep=10$) | **63.92**($ep=20$) | 62.58($ep=20$) |
| SKIP, $w=5$ | 62.47($ep=10$) | 63.44($ep=10$) | 62.94($ep=15$) |
| random | 31.51 ($k=40$) | 32.64 ($k=40$) | 31.68 ($k=20$) |

### Supplementary D: GO molecular function

#### Malaria

GO class distribution, 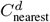 *Accuracy_GO_* for non-overlapping sequences and non-redundant sequences, for *Malaria*, are shown in S6, S7 and S8 respectively.

#### E.coli

GO class distribution, 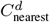 *Accuracy_GO_* for non-overlapping sequences and non-redundant sequences, for *E.coli*, are shown in S9, S10 and S11 respectively.

#### Yeast

GO class distribution, 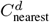 *Accuracy_GO_* for non-overlapping sequences and non-redundant sequences, for *Yeast*, are shown in S12, S13 and S14 respectively.

#### Human

The 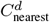 *Accuracy_GO_* for non-overlapping sequences, for *Human*, is shown in Table S16.

**Table S6.** GO molecular function distribution (Malaria)

| GO class | No. of domains |
| --- | --- |
| Catalytic activity | 676 |
| Binding | 440 |
| Structural molecule activity | 171 |
| Transporter activity | 63 |
| Molecular function regulator | 22 |
| Transcription regulator activity | 13 |
| Cargo receptor activity | 1 |
| Molecular carrier activity | 1 |

**Table S7.** Average 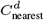 *Accuracy_GO_* for non-overlapping sequences, whenever *k* is not shown *k*=2, best shown in bold case (Malaria)

| Emb. Meth, $w$ | $dim=50$ | $dim=100$ | $dim=200$ |
| --- | --- | --- | --- |
| CBOW, $w=2$ | 58.2 ($k=5, ep=35$) | 58.24 ($k=5, ep=5$) | 57.87 ($ep=5$) |
| CBOW, $w=5$ | 57.87 ($ep=10$) | 56.94 ($ep=10$) | 57.1 ($ep=5$) |
| SKIP, $w=2$ | 59.48 ($k=5, ep=10$) | 59.82 ($ep=15$) | 58.68 ($ep=10$) |
| SKIP, $w=5$ | **60.61** ($ep=10$) | 59.01 ($ep=10$) | 59.39 ($ep=5$) |
| random | 46.21 ($k=40$) | 46.62 ($k=40$) | 45.99 ($k=40$) |

**Table S8.** Average 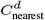 *Accuracy_GO_* for non-redundant sequences (*k*=2), best shown in bold case (Malaria)

| Emb. Meth, $w$ | $dim=50$ | $dim=100$ | $dim=200$ |
| --- | --- | --- | --- |
| CBOW, $w=2$ | 67.33 ($ep=5$) | 64.88 ($ep=5$) | 61.13 ($ep=5$) |
| CBOW, $w=5$ | 66.74 ($ep=5$) | 66.75 ($ep=5$) | 63.89 ($ep=5$) |
| SKIP, $w=2$ | 75.79 ($ep=35$) | 75.64 ($ep=45$) | 74.91 ($ep=5$) |
| SKIP, $w=5$ | 76.79 ($ep=10$) | **76.86** ($ep=40$) | 72.75 ($ep=20$) |
| random | 46.58 ($k=40$) | 46.22 ($k=40$) | 47.57 ($k=40$) |

**Table S9.** GO molecular function distribution (E.coli)

| GO class | No. of domains |
| --- | --- |
| Catalytic activity | 1565 |
| Binding | 476 |
| Transporter activity | 211 |
| Structural molecule activity | 117 |
| Transcription regulator activity | 39 |
| Molecular function regulator | 15 |
| Molecular carrier activity | 3 |
| Translation regulator activity | 1 |
| Molecular transducer activity | 1 |

**Table S10.** Average 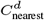 *Accuracy_GO_* for non-overlapping sequences, best shown in bold case (E.coli)

| Emb. Meth, $w$ | $dim=50$ | $dim=100$ | $dim=200$ |
| --- | --- | --- | --- |
| CBOW, $w=2$ | 67.66 ($k=5, ep=30$) | 67.46 ($k=20, ep=5$) | 67.34 ($k=20, ep=5$) |
| CBOW, $w=5$ | 67.78 ($k=5, ep=5$) | 67.46 ($k=20, ep=5$) | 67.34 ($k=20, ep=5$) |
| SKIP, $w=2$ | 68.15 ($k=5, ep=5$) | 67.54 ($k=5, ep=5$) | 67.75 ($k=20, ep=5$) |
| SKIP, $w=5$ | **69.1** ($k=5, ep=5$) | 67.82 ($k=5, ep=5$) | 68.15 ($k=5, ep=5$) |
| random | 64.46 ($k=40$) | 64.46 ($k=40$) | 64.46 ($k=40$) |

**Table S11.** Average 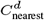 *Accuracy_GO_* for non-redundant domains (*k*=2), best shown in bold case (E.coli)

| Emb. Meth, $w$ | $dim=50$ | $dim=100$ | $dim=200$ |
| --- | --- | --- | --- |
| CBOW, $w=2$ | 71.41 ($k=5, ep=5$) | 68.45 ($k=5, ep=5$) | 67.87 ($ep=5$) |
| CBOW, $w=5$ | 74.95 ($ep=5$) | 71.69 ($ep=5$) | 68.91 ($ep=5$) |
| SKIP, $w=2$ | 81.27 ($ep=5$) | 80.32 ($ep=5$) | 80.36 ($ep=5$) |
| SKIP, $w=5$ | **81.72** ($ep=5$) | 81.64 ($ep=5$) | 80.77 ($ep=5$) |
| random | 64.38 ($k=40$) | 64.46 ($k=40$) | 64.38 ($k=40$) |

**Table S12.** GO molecular function distribution (S.cerevisiae)

| GO class | No. of domains |
| --- | --- |
| Catalytic activity | 1177 |
| Binding | 585 |
| Structural molecule activity | 208 |
| Transporter activity | 112 |
| Transcription regulator activity | 46 |
| Molecular function regulator | 40 |
| Translation regulator activity | 2 |
| Molecular transducer activity | 2 |
| Molecular carrier activity | 1 |
| Cargo adaptor activity | 1 |

**Table S13.**
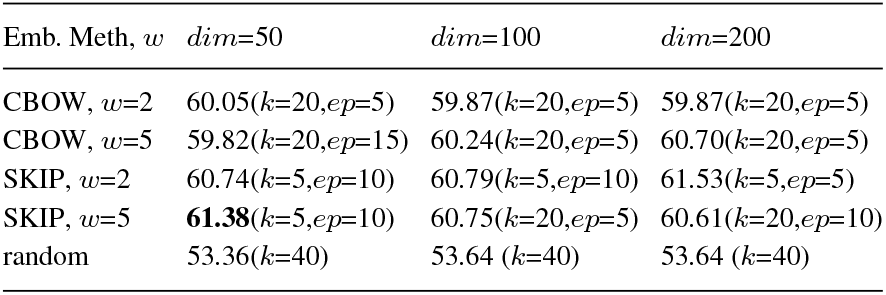
Average 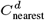 *Accuracy_GO_* fornon-overlapping sequences, best shown in bold (S.cerevisiae)

**Table S14.**
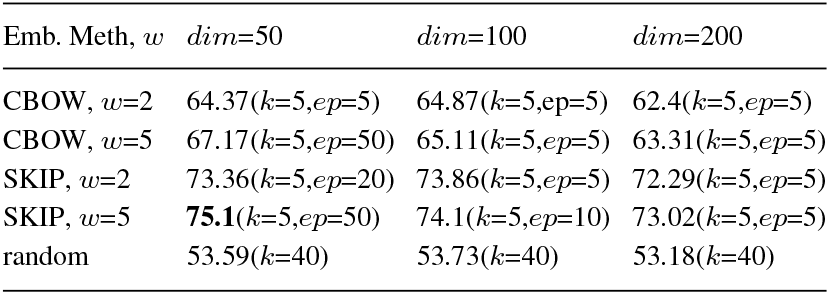
Average 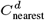 *Accuracy_GO_* for non-redundant sequences, best shown in bold (S.cerevisiae)

**Table S15.** GO molecular function distribution (Human)

| GO class | No. of domains |
| --- | --- |
| Catalytic activity | 1,945 |
| Binding | 1,583 |
| Transporter activity | 377 |
| Molecular transducer activity | 355 |
| Structural molecule activity | 262 |
| Transcription regulator activity | 203 |
| Molecular function regulator | 168 |
| Cargo receptor activity | 9 |
| Molecular carrier activity | 1 |
| Cargo adaptor activity | 1 |

**Table S16.** Average 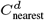 *Accuracy_GO_* for non-overlapping sequences, when *k* is not shown *k*=2, best shown in bold case (Human)

| Emb. Meth, $w$ | $dim=50$ | $dim=100$ | $dim=200$ |
| --- | --- | --- | --- |
| CBOW, $w=2$ | 57.7($ep=10$) | 58.82($ep=10$) | 58.04($k=5, ep=10$) |
| CBOW, $w=5$ | 58.1($ep=30$) | 58.9($ep=35$) | 58.2($ep=10$) |
| SKIP, $w=2$ | 60.51($k=5, ep=15$) | 60.67($ep=10$) | 59.3($ep=10$) |
| SKIP, $w=5$ | **60.59**($k=5, ep=35$) | 60.18($k=5, ep=10$) | 59.84($k=5, ep=10$) |
| random | 36.72 ($k=40$) | 36.44 ($k=40$) | 37.36 ($k=40$) |

### Supplementary E: Extrinsic evaluation

Class distribution for TargetP, Toxin and NEW data sets shown in Tables S17, S18 and S19 respectively. Model selection over hyperparameters, including architecture, shown in Table S20.

**Table S17.**
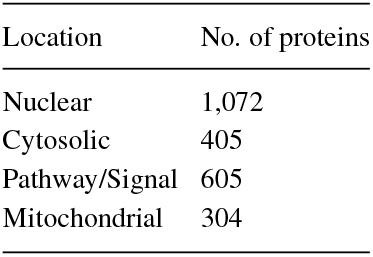
TargetP

**Table S18.**
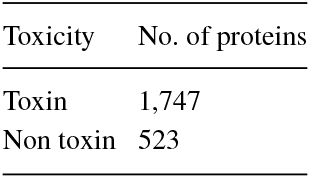
Toxin

**Table S19.** NEW

| EC Primary class | No. of proteins |
| --- | --- |
| Oxidoreductases | 2,234 |
| Transferases | 5,232 |
| Hydrolases | 4,099 |
| Lyases | 1,124 |
| Isomerases | 731 |
| Ligases | 1,124 |

**Table S20.** Average performance of simple neural architectures using as input dom2vec on inner three-fold cross validation. For Toxin AuROC is shown and for the rest of the data sets mc-AuROC is shown. Best values shown in bold case.

| Model \ Data set | TargetP | Toxin | NEW |
| --- | --- | --- | --- |
| CNN, size=(1,2),filters=200 | 0.9191 | 0.9074 | 0.9845 |
| CNN, size=1,filters=128 | **0.9288** | 0.8957 | 0.9844 |
| FastText (uni-gram) | 0.8829 | 0.9029 | 0.9818 |
| LSTM, dim=512,layer=1 | 0.9103 | 0.9025 | 0.9857 |
| bi-LSTM, dim=512,layer=1 | 0.921 | 0.9052 | 0.9857 |
| SeqVecNet dim=32 | 0.9017 | 0.9086 | **0.9876** |
| SeqVecNet dim=512 | 0.9206 | **0.9145** | 0.9864 |
| SeqVecNet dim=1024 | **0.9228** | 0.9034 | 0.9861 |

## Footnotes

1 In order to account for differences in specificity of different GO annotations, we always used the depth-1 ancestor of each annotation (that is, children of the root molecular function term, GO:0003674).

## Notes

### Competing Interest Statement

The authors have declared no competing interest.

### Summary of Updates

The changes are two-fold. Firstly, I have identify that in Toxin data set the proteins with unknown full-length domain is 46% of the data set so after removing them, I could pass the base lines, which use sequence embeddings so they didn't get affected in the first version of experiments. I have also inspected the issue of out-of-vocabulary, for domains as input features (equivalently words for NLP), for the TargetP data set. After having performed an experiment where the test had increased percentage of OOV I could show that dom2vec gets comparable results with the best performing embeddings (SeqVec) for a reasonable range of OOV. Greatly, I have changed the focus of the paper to reflect its potential. That is showing that unsupervised embedding methods, invented for words in natural language (e.g English), work also in protein domain in the "language" forms by domain architectures and capture context between collocated domains, supports the accepted theory of protein evolution by rearrangement and enrich the reported grammar of protein domain architectures. In a direct analogy of word embeddings capturing semantic and lexical similarity, protein domain embeddings, dom2vec, captures secondary structure, enzymatic function and GO molecular function similarity for protein domains in similar context. Consequently, we hope that these embeddings are not used only for performance gains, but also to answer biological questions on protein domain architectures based on data-driven experiments.

